# High-resolution 3D reconstruction of human oocytes using FIB-SEM

**DOI:** 10.1101/2021.03.30.437653

**Authors:** Zuzana Trebichalská, Jakub Javůrek, Martina Tatíčková, Drahomíra Kyjovská, Soňa Kloudová, Pavel Otevřel, Aleš Hampl, Zuzana Holubcová

## Abstract

The oocyte plays a pivotal role in the reproduction of our species. Nevertheless, its biology remains poorly understood. Electron microscopy is traditionally used to inspect the ultrastructure of female gametes. However, two-dimensional micrographs contain only fragmentary information about the spatial organization of the complex oocyte cytoplasm. Here, we employed the Focused Ion Beam Scanning Electron Microscopy (FIB-SEM) to explore human oocyte intracellular morphology in three dimensions (3D). Volume reconstruction from high-resolution image stacks provided an unprecedented view of ooplasmic architecture. Organelle distribution patterns observed in 9 donor oocytes, representing 3 maturational stages, documented structural changes underlying the process by which the egg acquires developmental competence. 3D image segmentation was performed to extract information about distinct organelle populations. The quantitative analysis of the organelle abundance revealed that mitochondrion occupies ~ 4.26 % of the maturing oocyte cytoplasm. This proof-of-concept study demonstrates the potential of FIB-SEM imaging to study human oocyte morphology.

## Introduction

The oocyte (egg) is the largest cell in the human body. When fertilized, exceptionally voluminous ooplasm constitutes the cellular mass of a newly-formed zygote. The female gamete’s influence over an embryo’s developmental fate is generally acknowledged [1], yet we still lack a clear understanding of human oocyte morphophysiology.

The transmission electron microscopy (TEM) proved to be a valuable tool for examining the intracellular morphology of reproductive cells at the nanoscale level [2]. However, the TEM micrographs are two-dimensional (2D) projections of a small area within the researched sample. Spatial attributes of biological structures pose a challenge to a single-plane image data interpretation and morphometric measurements. Volumetric imaging is required to elucidate how cells and tissues are organized in three dimensions (3D).

The 3D information can be retrieved by reconstruction of orderly stacked 2D image series. However, the acquisition of numerous serial sections, which must be manually handled, carefully registered, and precisely aligned, is arduous and error-prone.

Focused Ion Beam Scanning Electron Microscopy (FIB-SEM) represents an alternative to notoriously laborious TEM tomography. The dual-beam system automatically generates large sets of images with high resolution and an extensive field of view. This method relies on FIB-assisted ablation of a thin layer from the specimen’s block-face followed by SEM imaging of the newly exposed surface (Figure 1A). By repeating milling and scanning cycles, the desired 3D volume is recorded while the sample is progressively eroded. Unlike TEM, which harvests the power of a high energy electron beam passed through an ultrathin cross-section of the specimen, the SEM detects back-scattered and secondary electrons emitted from a limited depth beneath the ion-milled surface of the sample [3]. Traditional FIB employs easily controlled gallium (Ga) ions for high precision material removal. Given the relatively low ion currents (≤ 100nA), Ga FIBs are typically used for small samples not exceeding a few microns. Recently, FIB-SEM with xenon (Xe) plasma ion source gained popularity for its ability to achieve high ion currents (≤ 3μA) capable of large-scale milling tasks. The higher ablation rate offered by Xe FIB makes imaging of large volumes more time-efficient [4].

**Figure 1:**
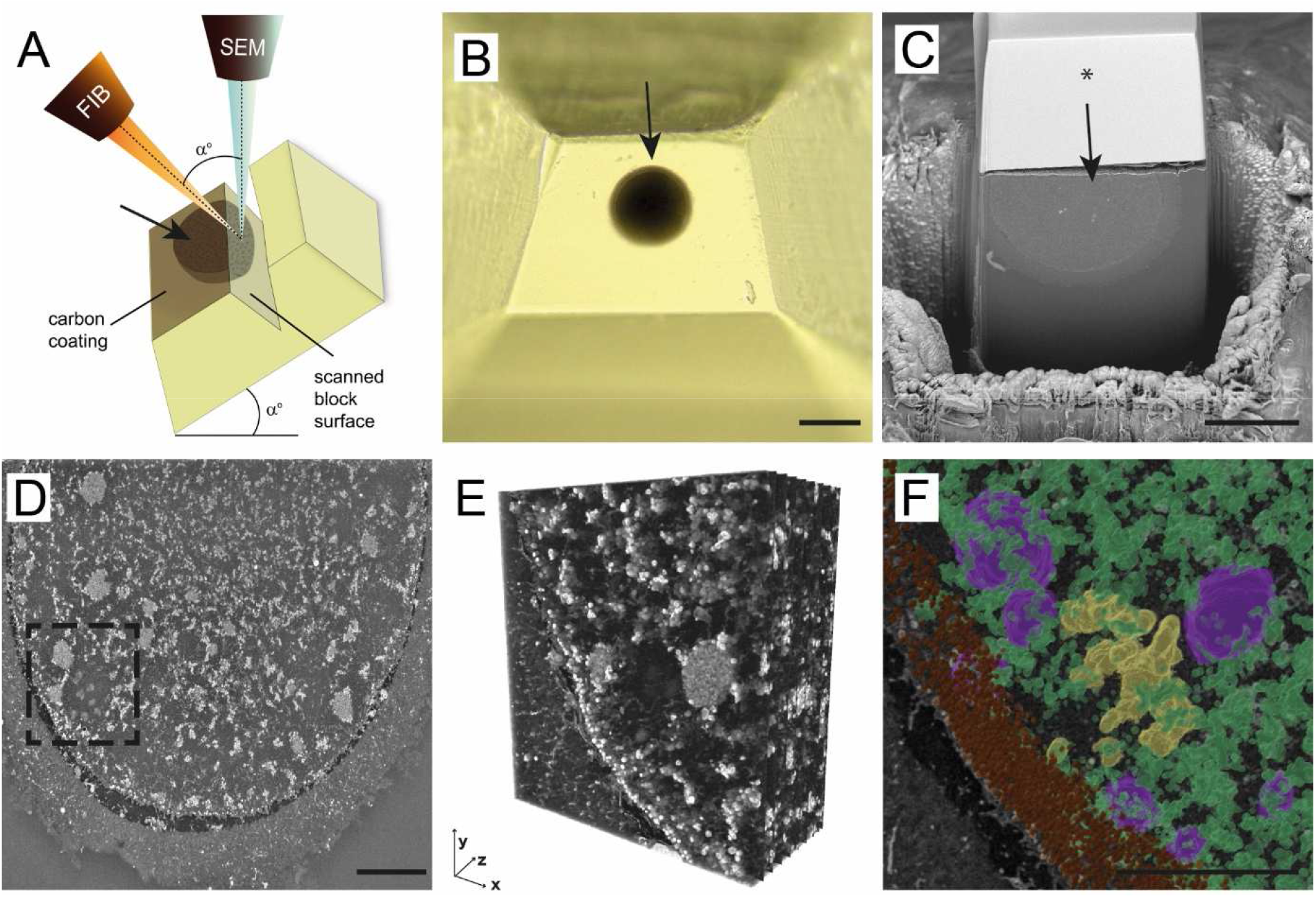
Overview of FIB-SEM imaging procedure and data processing. (A) The scheme of FIB-SEM setup. Both focused ion beam (FIB) and SEM (scanning electron microscope) columns are positioned in the microscope chamber, aiming at the scanned block surface’s coincidence point. The stage tilt corresponds to angle (a) between FIB and SEM column. The top of the surface of the resin block is coated with a carbon layer. Oocyte position is indicated with an arrow. (b) The top view of trimmed resin block with the sample embedded (arrow) (C) The trenched resin block with half of the oocyte (arrow). The block’s surface was coated with a thin carbon layer, and a silicon mask was placed over the sample (*). (D) The single grayscale image of MII oocyte surface scanned using FIB-SEM. (E) The 3D reconstruction of the spindle area (dashed rectangle in D). (F) The 3D segmentation of chromosomes (yellow), mitochondria (green), tubular endoplasmic reticulum (magenta), and cortical granules (brown). Scale bar: 100 μm (B), 50 μm (C), 10 μm (D, F).

Although initially developed for material sciences, FIB-SEM was applied to various biological specimens [5]. However, no study tested this technique’s feasibility to image the inner structure of large mammalian oocytes. Here, we report the first use of FIB-SEM tomography to visualize the 3D morphology of human oocytes. Using the automated slicing-scanning approach, we were able to reconstruct substantial volumes of ooplasm, examine spatial relations of microstructures, and quantify organelle population in rare samples of female sex cells.

## Results

A total of 9 human oocytes (3 germinal vesicle (GV), 3 metaphase I (MI), and 3 metaphase (MII) oocytes) were used to optimize conditions for large volume electron microscopy. The samples were processed using routine TEM protocol [6]. Instead of manual ultramicrotome sectioning, whole oocyte-containing resin blocks were individually inserted into the microscope chamber and subjected to FIB-SEM imaging (Figure 1A). In some experiments, the sample’s top part was removed to expose the cell’s central segment, thus reducing the image acquisition time (Figure 1B,C). Pre-conditioning of the sample surface with carbon coating was used to prevent excessive charging. We tested both Xe and Ga FIB-SEM systems and optimized milling and scanning conditions (technical parameters of each experiment are detailed in Supplementary Table I). A high current Xe beam sped up the trenching step, but ion currents >300nA exposed delicate samples to considerable heat stress. Automated Ga ion source reheating allowed us to run tomography without interruption, and currents up to 100 nA proved sufficient to image large volumes in a reasonable time. Additional functions such as automated drift correction and autofocus made the acquisition process robust and reliable. Hence, following appropriate optimization, both Xe- and Ga-based FIB-SEM systems are suitable for large volume microscopy of resin-embedded biological samples.

In our experiments, the continuous processing of bulk specimens lasted 5-90 hours, depending on the targeted volume and operating conditions. Improvement of imaging settings allowed us to visualize up to ~ 65 % of the human oocyte volume with 40-100 nm z-resolution (Supplementary Table I). The 3D reconstruction of the grayscale image stacks (120-1294 slices) provided a new perspective on female gametes’ morphology (Figure 1D-F; Video 1,2). In line with previous TEM studies [2, 6], the most prominent ultrastructural features we detected involved clusters of small rounded mitochondria, aggregates of the endoplasmic reticulum (ER), and cortical granules (Figure 1D,E). In MI and MII oocytes, chromosomes and microtubule bundles could be recognized in the meiotic spindle area enclosed by organelle-rich cytoplasm (Figure 1D-F; Videos 1,2). However, fine morphological details such as mitochondrial cristae or small vesicles were not discernible with the current settings and resolution used. Interestingly, all but one sample exhibited prominent electron-dense specs with a diameter reaching up to 7 μm (Figure 2C,F,I; Video 1). These bodies, known as cytoplasmic inclusions, were previously identified as tertiary lysosomes containing degraded intracellular material [6]. Although our FIB-SEM images did not achieve the level of detail seen in TEM micrographs, their 3D reconstruction provided an opportunity to analyze oocyte organelles’ spatial interaction (Video 1).

**Figure 2:**
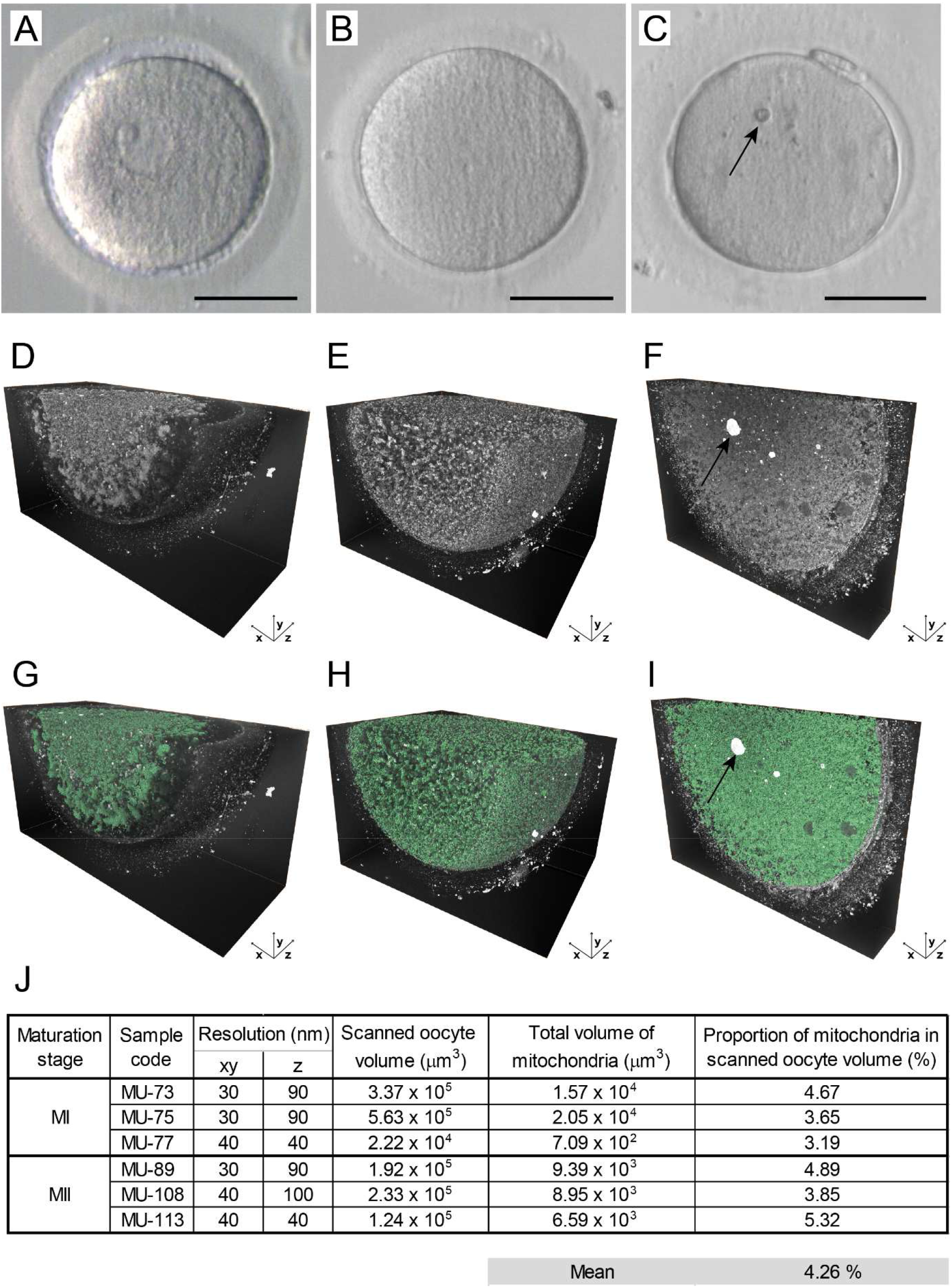
Large volume electron microscopy of maturing oocytes. (A-C) The live appearance of representative (A) germinal vesicle (GV), (B) metaphase I (MI), and (C) metaphase II (MII) oocyte in transmitted light. An arrow indicates prominent cytoplasmic inclusion. Scale bar: 50 μm (D-F) The histogram-enhanced 3D image of (D) GV, (E) MI, and (F) MII oocyte. An arrow indicates prominent cytoplasmic inclusion. (G-I) The 3D image in (G) GV, (H) MI, and (I) MII oocyte with mitochondria segmented (green). An arrow indicates prominent cytoplasmic inclusion. (J) Quantification of a total mitochondrial volume in analyzed MI and MII oocyte samples.

Comparison of 3D datasets obtained from GV, MI, and MII oocytes confirmed previous TEM findings that oocyte maturation is accompanied by rearrangement of cytoplasmic components. While the cortical area of prophase-arrested oocytes was devoid of organelles structures, in maturing oocytes, the detectable intracellular structures were evenly distributed throughout the cell mass (Figure 2A–I). Due to insufficient contrast, we could not observe the maturation-induced association of mitochondria and small ER cisternae documented by TEM. However, the large aggregates of tubular ER surrounded by a few mitochondria were detected in most MI and MII oocytes (Video 1, Video 2).

Software-enhanced 3D image segmentation was instrumental in dissecting spatial relationships among different structural elements and evaluating their prevalence and distribution within the cell (Figure 1F, Figure 2G–I, Video 2). For precise quantification, we focused on the most abundant organelles, the mitochondria, that exhibited typical morphology and sufficient contrast in all datasets (Figure 2G–I). The quantitative analysis of the 3D image data indicated that, in our samples of MI and MII oocytes, mitochondrion occupies approximately 4.26 % cytoplasm volume (Figure 2J). This result provides a more accurate estimate than previous 2D stereological studies [7, 8].

In summary, our work illustrates that volumetric imaging combined with advanced image analysis increase the yield of morphological data from precious biological samples, such as human oocytes.

## Discussion

To the best of our knowledge, this is the first study investigating the ultrastructure of human oocytes in 3D. Elucidating details about the spatial organization of complex ooplasm in developing female gametes improves our understanding of cellular processes conferring the egg with fertilization potential. This knowledge is vital for developing experimental and therapeutical strategies involving oocyte microsurgery, such as mitochondrial replacement therapy [9].

The procurement of human female gametes for research is limited to small numbers. It is thus crucial to maximize the amount of information obtained from a single human oocyte. Here, we used large volume electron microscopy to analyze the 3D ultrastructure of reproductive cells retrieved from young women with no history of fertility issues. Although the impact of ovarian stimulation and in vitro maturation on egg quality cannot be excluded, these samples are more representative for normal oocyte development than typically researched failed-to-be-fertilized eggs derived from patients struggling with infertility.

The volumetric image data presented here complemented our findings from the previous ultrastructural study [6]. Using the same fixation and sample processing protocol, we could directly compare attributes of our TEM micrographs and FIB-SEM images. While TEM represents a gold standard for scrutiny of minute morphological details, the extensive field of view of precisely sectioned FIB-SEM images is convenient for assessing the global organization of oocyte cytoplasm. Moreover, FIB-SEM instruments provide flexibility to switch from large-volume tomography to high-resolution imaging in the course of an experiment. Thus, selected regions of interest can be imaged with nanometer resolution.

Besides a wide range of magnifications, in situ manipulation and full automation represent the FIB-SEM technique’s main advantages. The common artifacts introduced during manual sectioning, such as folds, scratches, and contaminations, are avoided. However, considering this method’s irreversibly destructive nature, imaging protocol optimizations are required to minimize the risk of sample drifts, thermic distortion, excessive charging, and block-face curtaining, especially when working with rare samples. The technological advancements are aimed to balance the trade-off between resolution and speed of imaging. Future developments in precision instrumentation, big data management tools, and artificial intelligence-assisted software solutions promise to enhance FIB-SEM imaging’s time-efficiency of and facilitate 3D image data analysis.

In conclusion, this pioneering study paves the way for applying the FIB-SEM microscopy in human oocyte research. We believe that the prospect of visualizing whole oocyte volume in great detail would stimulate further studies looking into the inner workings of the cell our life starts with.

## Materials and methods

### Experimental material

The analysis was performed on 9 spare unfertilized oocytes derived from 9 young and healthy egg donors (aged 22-29 years, average age 25.56 years) who participated in a clinical egg donation program between February 2018 and November 2019. The donors’ mature eggs were utilized for fertility treatment, and surplus immature oocytes were used for research purposes, provided that written informed consent was obtained. The research study was undertaken under ethical approval issued by the Ethics Committees of collaborating institutions.

Ovarian stimulation and oocyte collection were carried out as previously described [6]. Immature oocytes were incubated in vitro until they reached the defined developmental stage. Each oocyte’s meiotic status was determined based on the presence/absence of a germinal vesicle (GV), the first polar body (PB), and metaphase I (MI)/metaphase II (MII) spindle detectable by polarized-light microscopy (PLM) [10]. Oocytes featuring an intact prophase nucleus (GV oocytes) were fixed no later than 3 hours after retrieval. Oocytes that showed a PLM-detectable spindle but no PB 3-6 hours of in vitro incubation were assigned to the MI oocyte category. Presence of a PB together with an MII spindle signal on the day after retrieval was considered a hallmark of MII oocyte maturity. Only normally appearing oocytes with no signs of dysmorphism were included in the study.

### Electron microscopy

The oocytes were fixed overnight in 3% glutaraldehyde (AGR1012, Agar Scientific, Stansted, UK) in 0.1 M sodium cacodylate buffer (C0250, Sigma Aldrich, St. Louis, USA) (pH 7.2–7.8) supplemented with 1% tannic acid (W304204, Sigma Aldrich, St. Louis, USA) at room temperature. Following post-fixation with 1% osmium tetroxide (O5500, Sigma Aldrich, St. Louis, USA) and 1.5% potassium ferrocyanide (P3289, Sigma Aldrich, St. Louis, USA) for 1 hour, the cells were individually embedded in 3% agarose (Sigma Aldrich, St. Louis, USA), dehydrated and embedded in epoxy resin (44611-4, Durcupan, Sigma Aldrich, St. Louis, USA).

Each block of hardened resin was manually trimmed into a pyramid shape with an oocyte positioned at its top (Figure 1B). After being coated with a 15 nm thin carbon layer, individual samples were inserted into a FIB-SEM equipped with an integrated Xe/Ga ion source (Tescan Amber (X)/Solaris X; Tescan Brno s.r.o., Brno, Czech Republic). The microscope stage was tilted to 55 degrees, corresponding to the angle between FIB and SEM columns (Figure 1A). Before starting image acquisition, the area around the sample was trenched, and the exposed block surface was smoothed out by FIB (Figure 1C). In order to reduce curtaining artifacts during FIB milling, a platinum protective layer was deposited on the surface of samples analyzed with Ga FIB, whereas silicon masks protected samples analyzed by Xe FIB (TESCAN TRUE-X sectioning method, described in detail in [11]). The FIB was operated at the energy of 30 keV and a beam current of 300 nA (Xe source) or 50-85 nA (Ga source) for trench etching and 100 nA (Xe source) or 20 nA (Ga source) for slicing and surface polishing. The SEM scanning was performed with accelerating voltage 3-5 kV, electron beam current 300-600 pA, working distance 5-6 mm, dwell time 10-32 us/px, and pixel size 30-40 nm (xy) / 40-100 nm (z).

### Image analysis

The generated image stacks were filtered using a 3D median in Fiji software [12]. For post-processing, the image stacks were imported into AMIRA software (Thermo Fisher Scientific, Waltham, Massachusetts, United States). The pixels that represented the mitochondria were automatically selected using the adjusted threshold option, and the individual z-sections were proofread and annotated to ensure the accuracy of automated detection. The other microstructures were segmented manually based on similar characteristics as shape, texture and signal intensity. The GV oocytes were excluded from the mitochondrial volume analysis because of the typically non-homogenous distribution of organelles. To generate animations, the segmented binary masks were converted into 3D models using Generate surface module. The scanned oocyte volume and total mitochondrion volume were quantified by Amira software based on the segmented data. The percentage of mitochondria was calculated per imaged oocyte volume.

## Acknowledgments

The authors acknowledge the Tescan Orsay Holding for expert support and access to FIB-SEM systems. We also thank the staff of Reprofit International for the recruitment of egg donors and the administration of informed consents. The work was funded by the Grant Agency of the Czech Republic (GJ19-14990Y).

## Competing interests

The authors declare no potential conflicts of interest concerning this article.

## Video 1

**3D reconstruction of the FIB-SEM-acquired image stack.** FIB-SEM-acquired stack of grayscale(-inverted) images and 3D reconstructions of MI/MII oocyte volume. Arrow indicates MI spindle with individual microtubules visible.

## Video 2

**3D visualization of distinct organelle populations in maturing oocytes**. Organelle segmentation in maturing oocytes. Representative examples of GV, MI, and MII oocytes: mitochondria (green), smooth ER (magenta), chromosomes (yellow), and cortical granules (brown).

## Supplementary Table

The imaging conditions of individual human oocyte samples.

| maturation stage | sample ID | FIB-SEM microscope | imaging time (h) | FIB settings |  |  | SEM settings |  |  |  | Dataset characteristics |  |  |  |  |  |
| --- | --- | --- | --- | --- | --- | --- | --- | --- | --- | --- | --- | --- | --- | --- | --- | --- |
|  |  |  |  | FIB type | trench milling (nA) | slicing (nA) | Detector type | Voltage (kV) | Beam current (pA) | Dwell time (µs/px) | Number of slices | Pixel XY (nm) | Pixel Z (nm) | XY pixel size | XY area (nm) | Z area (nm) |
| GV | MU-59 | Amber X | 39,8 | Xe | 300 | 100 | LE-BSE | 5 | 300 | 32 | 322 | 30 | 90 | 3165x1452 | 94950x43650 | 28980 |
|  | MU-107 | Solaris X | 43,5 | Xe | 300 | 100 | LE-BSE | 4 | 300 | 32 | 222 | 40 | 100 | 4224x1674 | 168960x66960 | 22200 |
|  | MU-158 | Amber | 49,4 | Ga | 50-85 | 20 | LE-BSE | 5 | 500 | 10 | 636 | 30 | 90 | 4899x2394 | 146970x71820 | 57240 |
| MI | MU-73 | Amber | 89,8 | Ga | 50-85 | 20 | LE-BSE | 5 | 600 | 10 | 1294 | 30 | 90 | 4124x2228 | 123720x66840 | 116460 |
|  | MU-75 | Amber | 76,2 | Ga | 50-85 | 20 | LE-BSE | 5 | 300 | 10 | 1125 | 30 | 90 | 4416x2622 | 132480x78660 | 101250 |
|  | MU-77 | Solaris X | 5,3 | Xe | 300 | 100 | LE-BSE | 4 | 300 | 32 | 120 | 40 | 100 | 1364x1462 | 54560x58480 | 12000 |
| MII | MU-89 | Amber X | 75,3 | Xe | 300 | 100 | LE-BSE | 3 | 400 | 30 | 237 | 30 | 90 | 5043x3315 | 151290x99450 | 21330 |
|  | MU-108 | Amber X | 22,7 | Xe | 300 | 100 | Axial | 5 | 300 | 32 | 451 | 40 | 100 | 2461x1715 | 98440x68600 | 45100 |
|  | MU-113 | Amber X | 55,1 | Xe | 300 | 100 | Axial | 3 | 300 | 19,2 | 494 | 40 | 40 | 3100x2152 | 124000x86080 | 19760 |

## Notes

### Competing Interest Statement

The authors have declared no competing interest.

## References

1. Mtango NR, Potireddy S, Latham KE. Oocyte quality and maternal control of development. Int Rev Cell Mol Biol. 2008;268:223–90. DOI: 10.1016/S1937-6448(08)00807-1

2. Sathananthan AH. Ultrastructure of human gametes, fertilization and embryos in assisted reproduction: a personal survey. Micron. 2013;44:1–20.DOI: 10.1016/j.micron.2012.05.002

3. Xu CS, Hayworth KJ, Lu Z, Grob P, Hassan AM, García-Cerdán JG, Niyogi KK, Nogales E, Weinberg RJ, Hess HF. Enhanced FIB-SEM systems for large-volume 3D imaging. Elife. 2017;6. DOI: 10.7554/eLife.25916

4. Hrnčíř T, Lopour F, Zadražil M, Delobbe A, Salord O, Sudraud P, editors. Novel plasma FIB/SEM for high speed failure analysis and real time imaging of large volume removal. In ASM International, EDFAS Organizing Committee; ISTFA 2012: 38th International Symposium for Testing and Failure Analysis; Phoenix Convention Center, Phoenix, Arizona, USA; November 11-15, 2012; 620. https://doi.org/10.31399/asm.cp.istfa2012p0026

5. Kizilyaprak C, Stierhof YD, Humbel BM. Volume microscopy in biology: FIB-SEM tomography. Tissue Cell. 2019;57:123–8. DOI: 10.1016/j.tice.2018.09.006

6. Trebichalská Z, Kyjovská D, Kloudová S, Otevřel P, Hampl A, Holubcová Z. Cytoplasmic maturation in human oocytes: an ultrastructural study †. Biol Reprod. 2021;104(1):106–16. DOI: 10.1093/biolre/ioaa174

7. Pires-Luís AS, Rocha E, Bartosch C, Oliveira E, Silva J, Barros A, Sá R, Sousa M. A stereological study on organelle distribution in human oocytes at prophase I. Zygote. 2016;24(3):346–54. DOI: 10.1017/S0967199415000258

8. Coelho S, Pires-Luís AS, Oliveira E, Alves Â, Leal C, Cunha M, Barreiro M, Barros A, Sá R, Souse M. Stereological study of organelle distribution in human oocytes at metaphase I. Zygote. 2020;28(4):308–17. DOI: 10.1017/S0967199420000131

9. Darbandi S, Darbandi M, Khorram Khorshid HR, Sadeghi MR, Agarwal A, Sengupta P, Al-Hasani S, Akhondi MM. Ooplasmic transfer in human oocytes: efficacy and concerns in assisted reproduction. Reprod Biol Endocrinol. 2017;15(1):77. DOI: 10.1186/s12958-017-0292-z

10. Holubcová Z, Kyjovská D, Martonová M, Páralová D, Klenková T, Kloudová S. Human Egg Maturity Assessment and Its Clinical Application. J Vis Exp. 2019(150). DOI: 10.3791/60058

11. Hrnčíř T, Šikula M, Oboňa JV, Gounet P. How to Achieve Artifact-Free FIB Milling on Polyimide Packages. In ASM International, EDFAS Organizing Committee; ISTFA 2016: 42nd International Symposium for Testing and Failure Analysis; 2016; Fort Worth, Texas, USA; November 6-10, 2016; 630–4. https://doi.org/10.31399/asm.cp.istfa2016p0630

12. Schindelin J, Arganda-Carreras I, Frise E, Kaynig V, Longair M, Pietzsch T, Preibisch S, RuedenC, SaalfeldS, SchmidB, TinevezJ-Y, White DJ, Hartenstein V, Eliceiri K, Tomancak P, Cardona A. Fiji: an open-source platform for biological-image analysis. Nat Methods. 2012;9(7):676–82. DOI: 10.1038/nmeth.2019

